# Enhanced anti-amyloid effect of combined leptin and pioglitazone in APP/PS1 transgenic mice

**DOI:** 10.1101/2020.06.24.168518

**Authors:** Yao Liu, Kelsey A. Hanson, Graeme McCormack, Justin Dittmann, James C. Vickers, Carmen M. Fernandez-Martos, Anna E. King

## Abstract

**Background:** Alzheimer’s disease (AD) has challenged single-target therapeutic strategies, raising the possibility that combined therapies may offer a more effective treatment strategy.

**Objective:** There is substantial evidence for the efficacy of leptin (L) (neuroprotective hormone) and pioglitazone (P) (anti-inflammatory agent) as monotherapies in AD. We have previouly shown that combination treatment of L+P in APP/PS1 mice at the onset of pathology significantly improved memory and reduced brain Aβ levels relative to control mice. In this new study, we sought to replicate our previous findings in a new cohort of APP/PS1 mouse to further confirm whether the combined treatment of L+P is superior to each treatment individually.

**Methods:** We have re-evaluated the effects of L+P co-treatment in APP/PS1 mice using thioflavin-S staining, MOAβ immunolabeling and enzyme-linked immunosorbent assay (ELISA) to examine effects on Aβ levels and pathology, relative to animals that received L or P individually. To explore mechanism of regulation, we used Western blotting to examine the expression of the peroxisome-proliferator activated receptor γ (PPARγ), due to its potential role in the regulation of the inflammatory response.

**Results:** We demonstrated that combining L and P significantly enhances the anti-Aβ effect of L or P in the hippocampus of APP/PS1 mice. Western blot analysis indicated that Aβ reduction was accompanied by up-regulation of the PPARγ levels.

**Conclusion:** Our findings suggest that combining L and P significantly enhances the anti-Aβ effect of L or P in the hippocampus of APP/PS1 mice, and may be a potential new effective strategy for AD therapy.

## 1. INTRODUCTION

Alzheimer’s disease (AD) accounts for approximately 60-70% of dementia cases. AD is characterized by extracellular amyloid-beta (Aβ) deposits, intracellular neurofibrillary tangles [1] and clinical symptoms of cognitive impairment. As current evidence indicates that Aβ deposits begin to form in the brain long before the onset of AD symptoms [2-5], interventions specifically aimed at disaggregating existing Aβ plaques and reducing Aβ oligomers may constitute a useful approach to AD therapy. However, anti-Aβ agents have not yielded expected outcomes clinically, which may be due to the incomplete removal of pre-existing Aβ aggregates. In addition, none of the developing anti-Aβ single-target drugs survived phase III clinical trials [6], which indicate that single-target drugs are limited in the treatment of AD. Thus, there is an urgent need to re-examine the validity of the Aβ-targeting approaches and to develop new effective strategies.

Pioglitazone (P), an agonist of nuclear peroxisome-proliferator activated receptor γ (PPARγ), is a drug that is used to treat type-2 diabetes through modulation of metabolic pathways. Recent studies have shown that P might provide a protective effect on AD risk among individuals with type-2 diabetes [7], suggesting that drugs used to treat diabetes may also be efficacious in AD. In this regard, P has anti-inflammatory properties [6; 8], improves spatial memory in AD mouse models and AD patients [9] and has demonstrated reduced Aβ plaques in experimental models [6; 10; 11], through mechanisms related to the amyloid degradation [12; 13]. The effect of P on reducing soluble Aβ species in models of AD may also be partly mediated by PPARγ [12].

A range of studies at both the clinical and pre-clinical level indicate that disrupted leptin (L) signalling pathways are linked to the pathophysiology of AD [14-18]. Furthermore, accumulating *in vivo* and *in vitro* studies have suggested that L has notable effects on reducing Aβ production [19; 20] and enhancing neuroprotection [21-24]. In the brain, the binding of L to the neuronal receptor Ob-Rb activates janus-tyrosine kinase 2 (JAK2), resulting in the activation of the phosphatidylinositol-3-kinase/Akt (PI3K/Akt), JAK/signal transducers and activators of transcription (JAK/STAT), and 5′ adenosine monophosphate-activated protein kinase/sirtuin (AMPK/SIRT) pathways, potentially reducing Aβ production and increasing its clearance, and reducing tau phosphorylation [25].

We have previously reported that co-treatment of the anti-inflammatory P, and the neuroprotective L, significantly reduced hippocampal Aβ levels (soluble Aβ and fibrillary Aβ plaque burden) in a transgenic (Tg) mouse model of AD expresses human presenilin 1 (PS1) with deltaE9 mutation and a chimeric mouse/human Aβ precursor protein (APP) harboring the Swedish mutation (APPswe/PSEN1dE9) [21]. However, in our first study, the L and P drugs were not tested independently under the same experimental conditions. In addition, the failure of translation from animal studies has led to the scientific community increasing research on potential therapeutics in animals, including performing replication studies [26-28], prior to progressing to clinical studies. Therefore, in the current study, we sought to replicate our previous study in a new cohort of APP/PS1 mice using thioflavin-S staining, MOAβ immunolabeling and enzyme-linked immunosorbent assay (ELISA) to examine effects on Aβ levels and pathology, relative to animals that received L or P individually. In addition, we examined the protein expression of PPARγ, by using Western blotting, due to the potential role for PPARγ expressed in astrocytes and microglial cells in the regulation of the inflammatory response [8].

## 2. METHODS

### 2.1. Animals

Male APP/PS1 (APPswe/PSEN1dE9) Tg mice and age-matched wildtype (WT) littermates maintained on a C57BL/6 background [26] were used in this study. The maintenance and use of mice and all experimental procedures were approved by the Animal Ethics Committee of the University of Tasmania (A0013939 & A0016308), in accordance with the Australian Guidelines for the Care and Use of Animals for Scientific Purposes. All analyses were conducted by personnel blinded to the animal genotype.

### 2.2. Experimental Design

Male APP/PS1 mice and WT littermate controls, were treated with drugs at 28 weeks of age, a time when plaques are beginning to develop [27]. Animals were administered daily treatments of L (delivered IN) and/or P (delivered IP) or vehicle control for two weeks as previously described with modifications [21]. Treatment groups are summarized in Table 1. Mice were sacrificed at 40 weeks of age. Brain tissue was used for the analysis of Aβ plaque load and Western Blot analysis. (Fig. 1). Numbers of mice used in each analysis are summarized in Table 2.

**Table 1.** Different treatments on WT and Tg mice.

| Tg: APP/PS1 mice | WT: wild type mice |
| --- | --- |
| Group 1: IN Leptin + IP Pioglitazone (L+P) | Group 5: IN Leptin + IP Pioglitazone (L+P) |
| Group 2: IN Leptin + IP Vehicle (L) | Group 6: IN Leptin + IP Vehicle (L) |
| Group 3: IN Vehicle + IP Pioglitazone (P) | Group 7: IN Vehicle + IP Pioglitazone (P) |
| Group 4: IN Vehicle + IP Vehicle (VH) | Group 8: IN Vehicle + IP Vehicle (VH) |

**Abbreviations:** IN, Intranasal; IP, intraperitoneal; Tg, Transgenic; VH, vehicle; WT, wild type.

**Table 2.** Numbers of mice used in each analysis.

|  | Tg VH | Tg L | Tg P | Tg L+P | WT VH | WT L | WT P | WT L+P |
| --- | --- | --- | --- | --- | --- | --- | --- | --- |
| ThioS | 6 | 9 | 7 | 5 | - | - | - | - |
| MOAβ | 5 | 9 | 6 | 3 | - | - | - | - |
| ELISA | 3 | 5 | 5 | 3 | - | - | - | - |
| WB | 3 | 5 | 5 | 3 | 5 | 5 | 5 | 5 |

**Abbreviations:** WT, wild type; Tg, Transgenic (refer to APP/PS1 mice); ThioS, thioflavin-S; ELISA, enzyme-linked immunosorbent assay; WB, Western blotting.

**Fig. 1.**
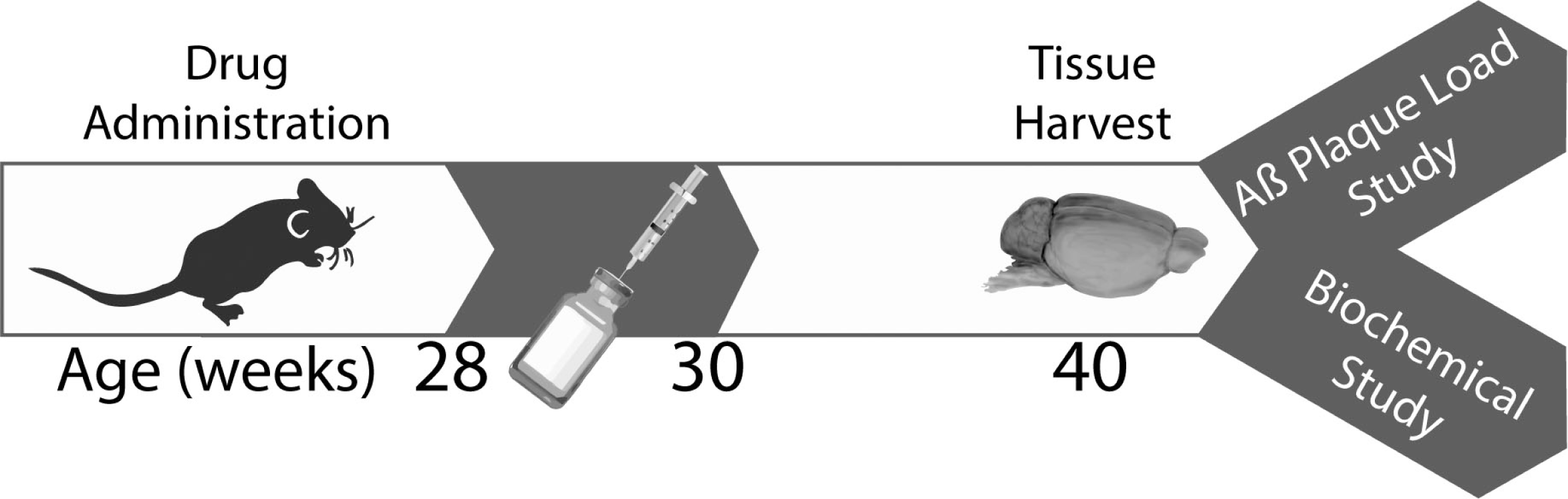
The schedule of experiment design. Male mice about 28 weeks underwent 2 weeks of drug administration. Ten weeks after the drug administration, the brain hemispheres were harvested for the analysis of Aβ plaque load and Western Blots.

### 2.3. Drug administration

Each genotype of mice was divided into four subgroups (n = 5**−**9 Tg mice/ subgroup and n = 10 WT mice/ subgroup) according to the treatments. Therefore, there were eight groups in total (Table 1). Pioglitazone (Takeda Pharmaceuticals) solution was prepared in 1:3 dimethyl sulfoxide (DMSO, Cayman Chemical): phosphate buffered saline (PBS, pH 7.2), and leptin solution was prepared in 0.125% (2.3mM) of N-tetradecyl-b-D-maltoside (TDM) (Sigma-Aldrich) reconstituted in PBS (pH 7.2) at a concentration of 1 mg/mL.

Mice received once daily IP injection of 150−200µL of 20mg/kg of pioglitazone solution or vehicle control (1:3 DMSO: PBS; pH 7.2). Following injection, IN treatments were given under light 3% isoflurane anesthesia. A 30µL pipette bearing a 20µL long gel loading tip was used to introduce 20µL of 0.03mg/kg leptin (Sigma-Aldrich) solution or vehicle control (0.125% TDM/PBS; pH 7.2) into the nasal cavity via only one of the external nares. Nares were alternated each day during the 2-week treatment starting with the left side. We selected IN administration for L delivery as it is a common technique in laboratory rodents, enabling administration of small drug volumes relative to other routes. Additionally, the nasal mucosa are richly supplied with blood vessels, potentially resulting in rapid substance absorption and subsequent systemic effects, avoiding the hepatic first-pass effect seen with oral delivery [28]. Both administration routes are commonly used as non-invasive drug administration techniques, which promote a minimal discomfort [28-30].

### 2.4. Tissue preparation

Animals were terminally anesthetized with sodium pentobarbitone (140 mg/kg) and transcardiacally perfused with 0.01M PBS (pH 7.4). Brains were immediately dissected, and right hemispheres were post-fixed overnight in 4% paraformaldehyde (PFA in 0.01M PBS), and then transferred to 18% and then 30% sucrose solutions overnight [31]. Serial coronal cryosections (40 µm thick) from right hemispheres were cut on a cryostat (Leica CM 1850). Tissue samples of neocortex (CTX) and hippocampus (HP) were dissected from left hemispheres and snap frozen in liquid nitrogen followed by storage at −80°C for later analysis.

### 2.5. Aβ plaque load

To determine Aβ plaque load in the CTX and HP, quantitation of fibrillar Aβ (Thioflavin-S staining) or total Aβ (MOAβ −2 immunolabeling) deposition was performed for four groups of APP/PS1 mice (Table 2). Serial coronal cryosections (40 µm thick) from right hemispheres were stained with thioflavin-S as previously described [32], to visualize fibrillar and dense-core Aβ plaques, or immunolabelled for Aβ plaques using the mouse anti-MOAβ-2 (1:1000; Novus Biologicals) antibody following antigen retrieval with formic acid (90%, 8 minutes) treatment [33], which specifically detects Aβ but not APP [34; 35]. Primary antibody binding was visualized using species specific fluorescent secondary antibodies conjugated to Alexa Fluor goat anti-mouse 488 (1:1000; Molecular Probes). Sections were mounted using fluorescent mounting medium (DAKO).

Slides containing brain sections were scanned via a VS120 slide scanner and whole brain sections imaged on a fluorescence microscope (Olympus). Images were analyzed with ImageJ software to quantify numbers and pixel areas of Aβ plaques. The whole CTX from the mid-line to the rhinal fissure from one hemisphere of the brain as well as the whole hippocampal formation was quantitated for each brain section. Segmentation of images was performed using the ImageSURF plugin which automatically segments images as plaques or background pixels by random forest classification [36]. Aβ plaque load (defined as percentage area thioflavin-S positive or MOAβ-2 labelling of the total area analyzed) was quantitated [33; 37]. In all cases, the specificity of immunoreactivity was confirmed by processing sections lacking primary antibody. All analyses were conducted by personnel blinded to animal genotype and treatment.

### 2.6. Protein preparation

The left sides of the CTX and HP from the four subgroups of WT and Tg mice (Table 2), respectively, were homogenized in RIPA buffer (Sigma) containing a protease inhibitor cocktail (Roche) and phosphatase inhibitors (AG Scientific) as previously described [38]. The samples were sonicated for 5 minutes and then centrifuged for 5 minutes at 13,000 RPM. The resulting supernatant was removed and stored at −20°C for Western blots or ELISA analysis. Protein concentration was determined with a bicinchoninic acid (BCA) assay kit (Invitrogen) according to the manufacturer’s instructions. Bovine serum albumin (BSA) was used as a standard.

### 2.7. Western blotting

Western blotting was performed as previously described [38] with modifications. Protein samples from the HP of all eight groups of mice (Table 2) were used. Proteins (15µg) were separated on 4-20% Bolt Bis-Tris Plus gels (Invitrogen), as described previously [15, 25]. The gel was subsequently transferred onto a polyvinylidene fluoride (PVDF) membrane (Bio-Rad) and blocked with 5% (w/v) non-fat milk tris-buffered saline (TBS) with 0.1% Tween-20 (Bio-Rad) for 1 hour at room temperature. The membrane was then incubated with rabbit anti-PPARγ (1:1000, Cell Signaling) primary antibody at 4°C overnight. Subsequently, a corresponding anti-rabbit horseradish peroxidase (HRP)-conjugated secondary antibody (1:5000; DAKO) was applied. Binding was visualized with enhanced chemiluminescence solution using Luminata Forte Western horseradish peroxidase (HRP) substrate (Millipore). Band intensity was measured as the sum optical density by using ImageJ software (version1.4; NIH) and normalized to control labelling of glyceraldehyde 3-phosphate dehydrogenase (GAPDH; 1:5000, Millipore).

### 2.8. Aβ_1-42_ ELISA

The quantitation of soluble human Aβ_1-42_ in hippocampus extracts was determined with a sandwich antibody ELISA kit (Invitrogen), according to the manufacturer’s instructions. Duplicate HP samples from four groups of APP/PS1 mice (Table 2) were used for ELISA analyses. Optical densities were read at 450nm on a microplate reader (Tecan), and concentrations of Aβ_1-42_ were determined by comparison to the standard curve using a 4-parameter algorithm. Soluble Aβ_1–42_ levels were normalized to total protein levels, and HP homogenates were expressed as picograms of Aβ_1–42_ content per milligram (pg/mg) of total protein [21].

### 2.9. Statistical analysis

Statistical analysis was performed using GraphPad Prism (version 6.0; GraphPad Software, La Jolla, CA, USA). All values are expressed as the mean ± standard error of the mean (SEM). Two-way analysis of variance (ANOVA) was used followed by Dunnett’s post hoc test, to compare all groups with control group (WT VH group or APP/PS1 VH group), while the non-parametric Student *t* test was used for the comparisons between APP/PS1 L and APP/PS1 L+P groups, and between APP/PS1 P and APP/PS1 L+P groups.

## 3. RESULTS

### 3.1. Combined treatment of L+P reduced Thioflavin-S–positive Aβ plaques in the hippocampus of APP/PS1 mice to a greater degree that either single treatment alone

To assess the Aβ deposition in the brain, we analyzed fibrillar Aβ plaque load by thioflavin-S staining (Fig. 2) and by immunostaining of total human oligomeric, and fibrillar forms of Aβ_1-42_ with the MOAβ-2 antibody (Fig. 3), in the HP and CTX of APP/PS1 L, APP/PS1 P, APP/PS1 L+P and APP/PS1 VH mice. Dunnett’s post hoc test demonstrated that L+P treatment significantly (p<0.05) reduced thioflavin-S–positive plaque deposition in the HP of APP/PS1 mice compared with APP/PS1 VH control group (Fig. 2A). In addition, APP/PS1 L mice as well as APP/PS1 P mice were not significantly different compared with APP/PS1 VH control group (Fig. 2A). These results indicated that the combination of L+P reduce thioflavin-S–positive Aβ plaque burden in the HP of APP/PS1 mice. However, there was no effect of any of the treatments on thioflavin-S–positive Aβ load in the CTX of APP/PS1 mice relative to APP/PS1 VH mice (Fig. 2B).

**Fig. 2.**
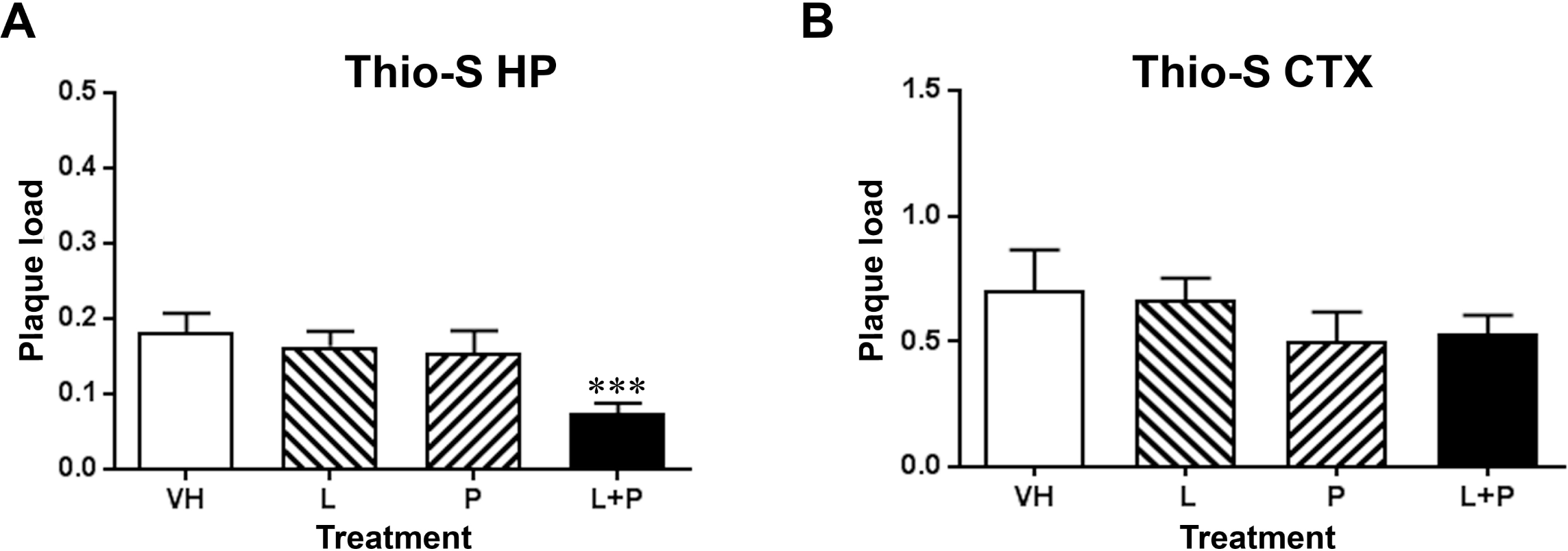
The analysis of thioflavin-S positive plaque load analysis. (a) Plaque load analysis showed the percentages of area stained by thioflavin-S were significantly (***p<0.01) lower in the HP of APP/PS1 L +P mice than in APP/PS1 VH mice. And there was no significant decrease of fibrillar Aβ plaques in HP of APP/PS1 APP/PS1 L or APP/PS1 P mice compared to APP/PS1 VH mice. (b) L alone or P alone or L and P therapy did not result in differences in fibrillar Aβ load in CTX of APP/PS1 mice relative to VH mice. Bar graphs represent the mean ± SEM. Statistical analyses were performed by two-way ANOVA followed by Dunnett test. Abbreviations: ANOVA, analysis of variance; CTX, neocortex; HP, hippocampus; L, leptin; P, pioglitazone; SEM, standard error of the mean; Tg, Transgenic (refer to APP/PS1 mice); VH, vehicle.

**Fig. 3.**
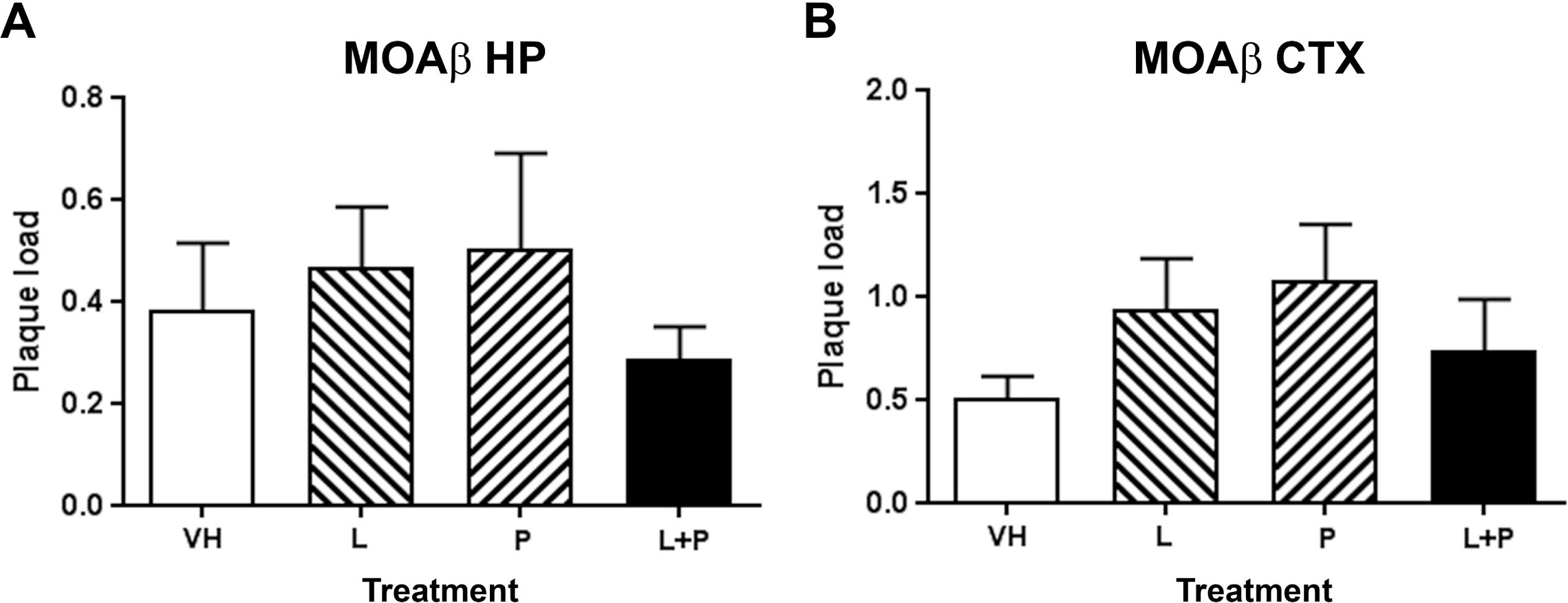
MOAβ-2–labelled Aβ load in the HP. There were no significant alterations to MOAβ-2–labelled Aβ load in either HP (a) or CTX (b) of APP/PS1 L, APP/PS1 P, APP/PS1 L +P mice relative to APP/PS1 mice. Bar graphs represent the mean ± SEM. Statistical analyses were performed by two-way ANOVA followed by Dunnett test. Abbreviations: ANOVA, analysis of variance; CTX, neocortex; HP, hippocampus; L, leptin; P, pioglitazone; SEM, standard error of the mean; Tg, Transgenic (refer to APP/PS1 mice); VH, vehicle.

Furthermore, there were no significant alterations to MOAβ-2–labelled Aβ load in either HP (Fig. 3A) or CTX (Fig. 3B) of APP/PS1 L, APP/PS1 P, APP/PS1 L+P mice relative to APP/PS1 VH mice.

### 3.2. Enhanced effect of combined treatment of L+P on levels of soluble Aβ_1-42_ in the hippocampus of APP/PS1 mice

We further performed sandwich ELISA using the extracts from the HP of four groups of APP/PS1 mice to determine the level of soluble Aβ_1-42_. Compared with APP/PS1 VH mice, the amounts of Aβ_1-42_ were significantly decreased in the mice administered with L alone (p<0.05) or L+P (p<0.01) (Fig. 4). In addition, the level of soluble Aβ_1-42_ decreased by 63.04% in the L-treated group and by 36.22% in the P-treated group, respectively. In the L+P group, Aβ_1-42_ levels dropped by 80.54% (Fig. 4).

**Fig. 4.**
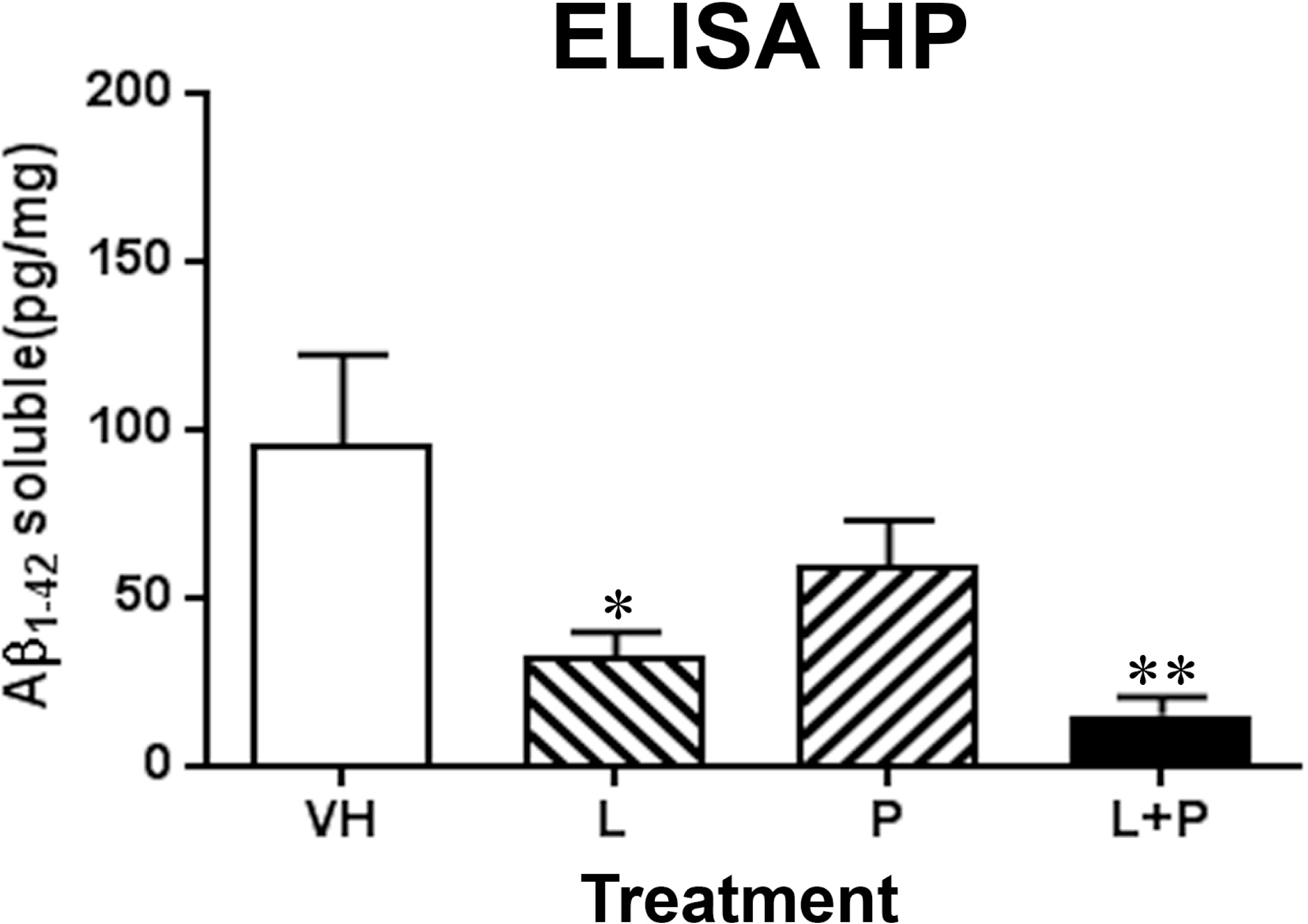
ELISA analysis of Aβ_1-42_ in the HP. The amounts of Aβ_1-42_ were significantly (*p<0.05, **p<0.01) decreased in the HP of APP/PS1 mice administered with L alone or L+P, compared with APP/PS1 VH mice. Bar graphs represent the mean ± SEM. Statistical analyses were performed by one-way ANOVA followed by Dunnett test. Abbreviations: ANOVA, analysis of variance, HP, hippocampus; L, leptin; P, pioglitazone; SEM, standard error of the mean; Tg, Transgenic (refer to APP/PS1 mice), VH, vehicle.

### 3.3. Effects of the treatments on the molecular regulation of PPARγ levels in the hippocampus of APP/PS1 mice

We next investigated if mechanism underlying the reduction of Aβ load in the brain of APP/PS1 mice could be partially mediated by the molecular regulation of PPARγ levels, which play a master role in the regulation of the neuroinflammatory processes in AD (Fig 5). Indeed, previous research has demonstrated that P reduces Aβ deposition in association with Akt/GSK3β activation by increasing the levels of insulin-degrading enzyme (IDE) and PPARγ expression [39]. For that reasons, in the current study, the downstream effect of treatment on PPARγ was investigated. Immunoblotting analysis showed PPARγ levels were significantly (P<0.05) increased in the HP of APP/PS1 VH mice relative to WT VH mice (Fig. 5A-B). Interestingly, PPARγ levels were significantly increased in WT mice following L+P treatment (Fig. 5B). There was also a significant (p<0.05) increase of PPARγ in the HP of APP/PS1 L mice, APP/PS1 P mice and APP/PS1 L+P mice, relative to APP/PS1 VH mice (Fig. 5B). Remarkably, combined treatment of L+P resulted in a significant (p<0.05) increase of PPARγ in the HP of APP/PS1 mice, relative to single treatment of L or P (Fig. 5B).

**Fig. 5.**
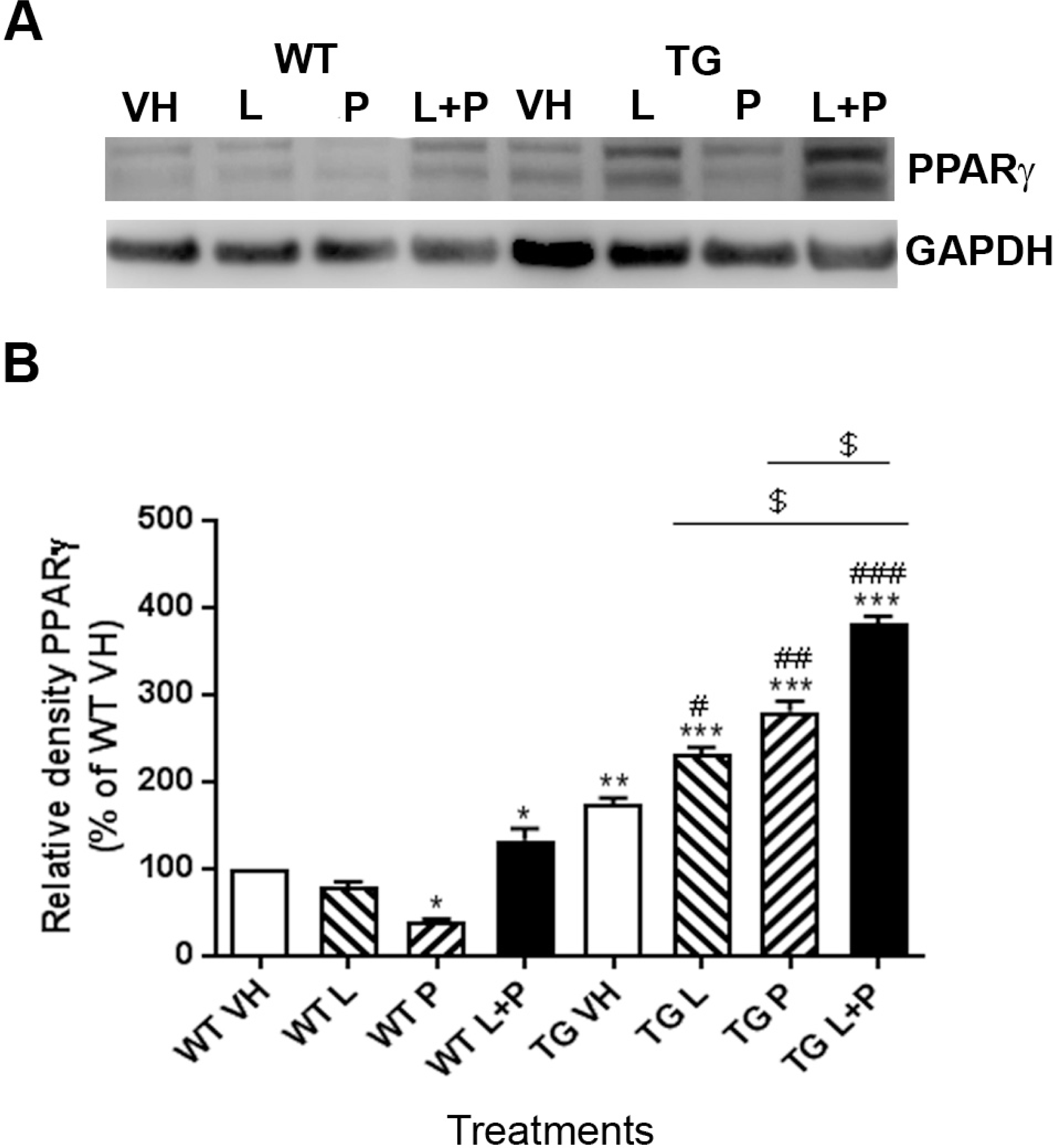
Immunoblotting analysis of the expression of PPARγ Protein. (a) Representative GAPDH-normalized immunoblotting images. (b) Quantitation of PPARγ in HP extracts from in the HP of WT and APP/PS1 mice administered with L alone, P alone or L+P, and APP/PS1 VH mice at 40 weeks. The combination of L and P increased the expression of PPARγ protein in APP/PS1 mice, compared with the single treatment of L or P. Data are expressed as a % of the WT VH group. Bar graphs represent the mean ± SEM. Statistical analyses were performed using two-way ANOVA followed by Dunnett test (*p < 0.05, **p<0.01, ***p<0.001 versus WT VH group; ^#^p< 0.05, ^##^p< 0.01, ^###^p< 0.001 versus APP/PS1 VH group), while the nonparametric Student *t* test was used for the comparisons between APP/PS1 L and APP/PS1 L+P groups, and between APP/PS1 P and APP/PS1 L+P groups (^$^p< 0.05, ^$$^p< 0.01, ^$$$^p< 0.001, *t* test). Abbreviations: ANOVA, analysis of variance; HP, hippocampus; L, leptin; Tg, Transgenic (refer to APP/PS1 mice); P, pioglitazone; SEM, standard error of the mean; VH, vehicle; WT, wild type.

## 4. DISCUSSION

Numerous drugs have been proposed for AD, however there have been no effective treatments developed to date. The majority of therapeutic agents have been developed as monotherapies however, studies suggest that some combination therapies may have synergistic effects in AD (e.g. memantine and donepezil combination therapy [40]. Thus, there is a great need for studies using combination therapy in AD, based on the available drugs. We have previouly shown that combination treatment of L+P in APP/PS1 mice at the onset of pathology significantly improved memory and reduced brain Aβ levels (soluble Aβ and Aβ plaque burden) relative to APP/PS1 VH mice [21]. Our present study confirmed these findings, and further show that L+P effects on plaque load and soluble Aβ_1-42_ were greater than either L alone or P alone, and that combination treatment further reduced soluble Aβ_1-42_. Therefore, taken together both studies further confirm L+P combination treatment might have additive or synergistic anti-Aβ effects, suggesting that this combined anti-Aβ therapy may be more effective than the monotherapy of L alone or P alone.

Aβ is currently a major target of AD therapy, as self-assembly of Aβ monomers into soluble oligomers and/or insoluble fibrils is linked to cognitive impairment of AD [41]. Therefore, many current therapeutic approaches are based on targeting the reduction of Aβ levels and preventing the production and aggregation of Aβ [42-45]. Failures of human clinical trials in anti-Aβ monotherapies and a growing awareness of the complexity of AD have led to a focus on the potential of combination treatments as a new approach to reduce AD pathology. For example, co-administration of 4-(2-hydroxyethyl)-1-piperazinepropane-sulphonic acid (EPPS), an amyloid-clearing chemical, and donepezil, an acetylcholinesterase inhibitor, led to a rapid and consistent cognitive improvement in AD Tg mice [46].

A new direction for AD research is identifying drugs already used clinically to treat other diseases and to potentially employ them in combination [47]. Indeed, a minimum of six different agent classes (including anti-inflammatory agents, e.g. pioglitazone) are approved for clinical use, or are ready for phase III clinical trials as monotherapy in AD [48; 49]. These therapeutic agents have a potential to be used as combination therapy but, only a few clinical studies have been published that have investigated the effects of a combination therapy versus monotherapy on AD [48]. Our current study suggests the combination treatment of the neuroprotective hormone (L), and the anti-inflammatory and antidiabetic drug (P), may have additive or synergistic effects on reducing brain Aβ levels through signalling pathways, highlighting the combined treatment of L and P as a promising treatment for AD. P has been shown to be effective and well-tolerated in the treatment of patients with type-2 diabetes [50]. A phase II study of P in AD showed that P is safe and well-tolerated [51]. In addtion, L-replacement therapy is an effective and safe treatment for long-term improvement of glucose and lipid metabolism and complications in generalized lipodystrophy [52]. Therefore, it is highly likely that combinatorial treatment of L+P may be a safe and effective therapy. And L+P treatment may be a successful translation from animal models to clinical trials for AD in the future.

In the current study, we examined a key downstream pathway of leptin and pioglitazone signalling, PPARγ. Our finding that PPARγ was increased in the HP of APP/PS1 mice following L or P alone, or L+P confirms reports demonstrating a significant increase in PPARγ protein levels in neurons incubated with P [53] and in spinal cord injury after L treatment [54]. PPAR γ agonists, including P, have been shown to induce neuroprotective effects from Aβ-peptide neurotoxicity on hippocampal neurons [55]. In cultured cells, PPARγ overexpression diminishes Aβ production [56]. Moreover, by inducing PPARγ expression and activation, L can reduce beta-secretase 1 (BACE1) expression [57]. These findings in the current study suggest that upregulation of PPARγ may be responsible for reduced Aβ load.

In summary, to our knowledge, our study is the first to demonstrate that combined treatment of L+P reduces soluble Aβ and fibrillary Aβ plaque burden in the hippocampus of APP/PS1 mice by a possible synergistic or additive mechanism of the individual drugs. Our findings indicated that this combination of the neuroprotective hormone and the anti-inflammatory therapeutic strategy would result in a further Aβ reduction than the single treatment of L alone or P alone. Although combinatorial treatment of L +P may be a safe and effective therapy, further pharmacoepidemiologic studies are still warranted before the clinical use of this combined treatment in AD.

Furthermore, we found that L+P increased the protein expression of PPARγ, which could potentially improve memory function via the transcriptional regulation of BDNF expression [58]. Indeed, it has been suggested that PPAR*γ* agonists prevent the impairment of synaptic plasticity by increasing BDNF expression and dendrite spine density [59]. However, future studies will be necessary to determine whether combining L+P treatment also improves learning through regulation of brain Aβ levels in AD. Our data indicate that this combination therapy may show promise in the clinic as a pathway to reducing Aβ burden.

## 5. ACKNOWLEDGEMENTS

The authors would like to gratefully acknowledge Jamuna Chhetri and Alexander Cronk for excellent technical support. The work was supported by the funding from the JO and JR Wicking Trust (<u>Equity Trustees</u>).

## 6. AUTHOR CONTRIBUTIONS

Conceptualization: Carmen M. Fernandez-Martos, James C. Vickers, and Anna E. King.

Data Analysis: Yao Liu, Kelsey A. Hanson and Carmen M. Fernandez-Martos.

Investigation: Yao Liu, Kelsey A. Hanson, Graeme McCormack, and Justin Dittmann.

Funding Acquisition: James C. Vickers and Anna E. King.

Methodology: Yao Liu, Carmen M. Fernandez-Martos, Anna E. King, and James C. Vickers.

Resources: James C. Vickers and Anna E. King.

Supervision: James C. Vickers, Carmen M. Fernandez-Martos and Anna E. King.

Writing-original draft: Yao Liu.

Writing-reviewed &editing: Anna E. King, Carmen M. Fernandez-Martos, Kelsey A. Hanson, James C. Vickers.

